# Novel Covariance-Based Neutrality Test of Time-Series Data Reveals Asymmetries in Ecological and Economic Systems

**DOI:** 10.1101/044495

**Authors:** Alex Washburne, Josh Burby, Daniel Lacker

## Abstract

Systems as diverse as the interacting species in a community, alleles at a genetic locus, and companies in a market are characterized by competition (over resources, space, capital, etc) and adaptation. Neutral theory, built around the hypothesis that individual performance is independent of group membership, has found utility across the disciplines of ecology, population genetics, and economics, both because of the success of the neutral hypothesis in predicting system properties and because deviations from these predictions provide information about the underlying dynamics. However, most tests of neutrality are weak, based on static system properties such as species-abundance distributions or the number of singletons in a sample. Time-series data provide a window onto a system’s dynamics, and should furnish tests of the neutral hypothesis that are more powerful to detect deviations from neutrality and more informative about to the type of competitive asymmetry that drives the deviation.

Here, we present a neutrality test for time-series data. We apply this test to several microbial time-series and financial time-series and find that most of these systems are not neutral. Our test isolates the covariance structure of neutral competition, thus facilitating further exploration of the nature of asymmetry in the covariance structure of competitive systems. Much like neutrality tests from population genetics that use relative abundance distributions have enabled researchers to scan entire genomes for genes under selection, we anticipate our time-series test will be useful for quick significance tests of neutrality across a range of ecological, economic, and sociological systems for which time-series data are available. Future work can use our test to categorize and compare the dynamic fingerprints of particular competitive asymmetries (frequency dependence, volatility smiles, etc) to improve forecasting and management of complex adaptive systems.

**Author Summary:** From fisheries and forestries to game parks and gut microbes, managing a community of organisms is much like managing a portfolio. Managers care about diversity, and calculations of risk - for extinction or financial ruin - require accurate models of the covariance between the parts of the portfolio.

To model the covariances in portfolios or communities, it helps to start simple with a null model assuming the equivalence of species or companies relative to one another (termed “neutrality”) and letting the data suggest otherwise. Researchers in biology and finance have independently entertained and tested neutral models, but the existing tests have used snapshots of communities or the variance of fluctuations of individual populations, whereas tests of the covariances between species can better inform the development of alternative models.

We develop a covariance-based neutrality test for time-series data and use it to show that the human microbiome, North American birds, and companies in the S&P 500 all have a similar deviation from neutrality. Understanding and incorporating this non-neutral covariance structure can yield more accurate alternative models of community dynamics which can improve our management of “portfolios” of multi-species systems.

## Introduction

*“(A)s more individuals are produced than can possibly survive, there must in 50 every case be a struggle for existence, either one individual with another of the 51 same species, or with the individuals of distinct species, or with the physical 52 conditions of life.”* - Charles Darwin, Origin of Species [1]

Adaptive evolution requires that rivalrous goods are consumed by agents, those agents have heritable variation in how they acquire and consume the rivalrous goods, and the fitness of agents increases with the amount of goods consumed [2]. By Lewontin’s listing of the necessary conditions of evolution, a variety of systems can be seen as evolving. Genes in populations, species in a community, companies in a market, and political groups in a society all satisfy Lewontin’s axioms [2].

Canopy space is a rivalrous resource in the multi-species closed-canopy forests. If nothing else intervenes, a competitively superior tree species will dominate the canopy just like a competitively superior gene will become fixed in a population. In economic systems, companies compete over capital, customers, and labor, and a company well-adapted to a market will increase its share of the resources. In social systems, political groups compete over votes and occupied positions of power, and political groups with superior recruitment compared to other groups - either by persuasion, coercion, aggression, or reproduction with vertical transmission of culture - will increase the votes it receives and/or its representation in various positions of power.

These generalized competitive systems are examples of “complex adaptive systems” [3, 4, 5] and understanding how they evolve can provide insight into the drivers of adaptive evolution [6], diversity maintenance in human and natural systems [7], portfolio construction in a market [8], and problems of recognition and representation in multicultural societies [9]. Much literature has explored the stochastic fluctuations of individual populations (e.g. [10, 11]) or asset prices [12] in these systems, and accurate models of the stochastic time-evolution of multi-species systems can enable calculations of the risk of extinction [13], the dynamics of diversity (such as the entropy or evenness of a system), portfolio analysis, and other features of interest.

A common stochastic model in which all groups are functionally equivalent, termed “Neutral Theory” in ecology and population genetics, has been used across many systems [14, 15, 16, 17, 18]. By “functionally equivalent”, we mean that every agent’s performance in acquiring the rivalrous resource is independent of their group membership. In other words, an organism’s species identity, a company’s strategy or sector, a citizen’s political identity, or a political party’s platform have no impact on their ability to hold or acquire new rivalrous resources. Neutrality is a parsimonious starting point for community modeling because it is based on first principles of random birth and death or acquisition and release of resources that are appropriate for many competitive systems, and, because neutrality does not assume particular traits that distinguish groups and complex interactions between groups, it is invariant to grouping: the populations of neutral species can be aggregated into larger groups whose competition is also neutral.

Neutrality is often posed as a null model for multi-species systems because it can be parsimonious to assume, initially, that all species are equivalent. The mathematical tractability of neutral systems has allowed for useful calculations [19] that can sometimes accurately describe features of the system. However, despite the mathematical ease, calculations for features such as extinction time or the dynamics of portfolio diversity based on neutrality may be inaccurate for systems with non-neutral dynamics such as positive or negative frequency-dependent selection. Thus, there is a need for powerful and informative tests of neutrality to assess whether or not the dynamics of the competitive system are neutral.

In population genetics, tests of neutrality [20, 21] have facilitated rapid conceptual and empirical advancements [22], allowing researchers to scan entire genomes for neutral loci and identify loci that have been under selection. Neutrality tests developed in ecological and sociological systems test features of rank-abundance and frequency-abundance distributions [16, 23, 24]. Many of these existing neutrality tests utilize snapshots of a competitive system, but time-series contain a tremendous amount of data and can enable stronger tests of neutrality.

Some work has been done developing and utilizing tools to test whether or not population dynamics are consistent with Neutral Theory [10, 25, 11]. These tests rely on a particular description of Neutral Theory as the one-step process [26] posed by Kimura and Hubbell, but the proof [27] that the nonzero-sum volatility-stabilized market models [17] converge to neutral drift in relative abundances motivates a broader definition and more general tests of neutrality. Since neutrality is the per capita equivalence between species, it is necessarily relative, not absolute; a population is not neutral per se, but can only be neutral relative to another population. To the best of our knowledge, the existing time-series tests have all analyzed whether or not the variance in jumps in abundance increase linearly or quadratically with the population size prior to the jump, and none have examined the covariance structure of fluctuations in relative abundance.

Here, we present, to our knowledge, the first covariance-based neutrality test for time-series data. Our test provides deeper insight into the nature of non-neutrality than traditional tests of rank-abundance distributions and the volatility of individual populations. Our test utilizes the grouping invariance of neutral systems to isolate and test the covariance between changes is species’ relative abundances over small time intervals, allowing a rejection of neutrality for the entire community considered. We apply our test to 6 metagenomic time-series [28], a time-series of breeding birds across North America [29], and a time-series of market capitalization of companies in the S&P 500 from 2000-2005. We show that even some systems whose rank-abundance distributions appear neutral can exhibit significantly non-neutral covariances between species detected by our test. Furthermore, our test, based on random groupings of species, illustrates how to analyze the volatility of randomly formed groups to reveal state-dependent volatility that differs from neutrality. The non-neutral state-dependent covariance structure uncovered here can be incorporated to improve our models of community dynamics and calculations of species’ extinction times, portfolio risk, and more.

Figure 1: Illustration of our method **(A)** The dynamics of a 15-species neutral community of 10,000 individuals and migration probability m = 0.0002 (shown here) can be approximated by a WFP with λ = 20. If the community is neutral, then the CVTs should yield homoskedastic plots of *v*_*t*_ versus *f*_*t*_- We test neutrality by randomly drawing from the 2^*n*^ possible CVTs, performing homoskedasticity tests on *v*_*t*_ versus *f*_*t*_, and then testing the uniformity of the resulting P-value distribution using a modified KS-test (see details in supplement section S3). (B) The relative abundances of 15 independent, mean-reverting geometric Brownian motions, *d* log *X*_*t*_ = *μ*(*b* — log *X*_*t*_)dt + *σdW*_*t*_ with *μ* = 15, *σ* = 30, *b* = 10. Neutrality is rejected by the highly non-uniform distribution of P-values. The left-skewed P-value distribution indicates many CVTs had volatilities that depended on the state variable, *f*_*t*_.

## Results

Our method is proven analytically in the Materials & Methods section, and a demonstration of the method is provided in Figure 1.

We apply our neutrality test to 8 different datasets. Six of these datasets are sequence-count data of microbial communities [28] from three body sites on two individuals. One dataset is survey of breeding birds across North America from 1966-2014 [29], and one dataset is financial data, obtained from the Center for Research in Security Prices, of the day-end market shares and market capitalization of 451 companies in the S&P 500 from January 1, 2000 to January 1, 2005. These datasets are long, time-series datasets, many of which have rank-abundance distributions that are decently fit by neutral theory’s expected rank-abundance distribution (see supplement part S1 for a detailed description of the datasets and fits of neutral theory’s rank-abundance distributions).

Figure 2: Applying our test to time-series datasets reveal non-neutral competitive dynamics in microbial and financial systems. Goodness of fit P-values displayed are from a modified KS-test which accounts for dependence among the observations. Despite decent fits of neutral species-abundance distributions, our time-series test reveals that competitive asymmetries are important drivers in all systems except the female tongue bacteria.

Our test relies on multiple groups of species. To group the species, we randomly selected *a*_*i*_ = ±1, for 4,000 independent groups, we then calculated *v*_*t*_ as in equation (6), and used a White test [30] to test the homoskedasticity of *v*_*t*_. The White test performed auxiliary regression with a generalized linear model with a log-normal link function of the form

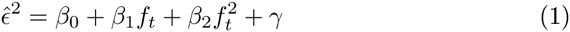

where 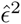 are the squared residuals from similar quadratic regression of *f*_*t*_ on *v*_*t*_. With a 0.005 significance threshold, neutrality was rejected in all but one dataset (figure 2).

Rejecting neutrality for these competitive systems motivates further investigation on whether the rejection of neutrality stems from sampling error or from true competitive asymmetries in the system. For the financial datasets, there is no sampling error - the reported values of day-end prices are the true values. For the metagenomic datasets, sampling error for sequence count data could be driving apparent non-neutrality.

Figure 3: **(A-D)** Analyzing the non-neutrality of competitive systems **(A)** The negative relationship between *v*_*t*_ vs. *f*_*t*_ indicates mean reversion. Overlaying *v*_*t*_ vs. *f*_*t*_ scatter plot from a particular CVT from the male tongue data onto the results from 4,000 WFP trajectories with long sampling intervals, Δt, shows that mean reversion can be accounted for by sparse time-sampling of the data. (B) However, even when correcting for sparse time-sampling, the left-skewed P-value distribution in the male tongue indicates stronger signal of non-neutral volatility than 16,000 surrogate WFPs. (C) The parameter *β*_2_ from significantly (*P* < 0.001 for male tongue, *P* < 0.01 for surrogate data) heteroskedastic auxiliary regressions in equation 7 reveals significantly more *β*_2_ > 0 than *β*_2_ < 0 in the data. The different P-value cutoffs are for visualization - the same bias for *β*_2_ > 0 holds for a standard cutoff of *P* < 0.01 (D) Overlaying scatterplots of the residuals, 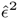, from all heteroskedastic cases (*P* < 0.01) of the male tongue data, reveals the empirical pattern of heteroskedasticity. Compared to surrogate neutral data, the male tongue is more volatile when the groupings are uneven, suggesting that either rare or abundant groups are more volatile - or equal groupings are relatively less volatile - than neutrality would predict. (E) All datasets have the same over-abundance of *β*_2_ > 0 for heteroskedastic (*P* < 0.05) CVTs.

Figure 3 examines the competitive asymmetry in the male tongue’s bacterial community. Scatter plots of *v*_*t*_ vs. *f*_*t*_ reveal a downward trend indicative of mean reversion - jumps in *f*_*t*_ are positive when *f*_*t*_ is above its mean and negative when *f*_*t*_ is above its mean (figure 3A). The mean reversion is regressed out prior to our auxiliary regression, but such strong mean reversion is not apparent in the simulated neutral community of figure 1. The discrepancy between the mean reversion in the data and the simulated neutral community may be due to a stronger and/or non-linear mean reversion in the microbial system, or it may be due to the relatively long time between time points in the data relative to the turnover rate of the community. Such sparse time-sampling could affect the accuracy of our test, which relies on the assumption that Δ*t* in equation 3 is small. A neutral community may still have mean reversion due to migration from a metacommunity or mutation/conversion rates between classes of agents, and sparsely sampling a time-series of such a neutral community may yield the same downward trend on plots of *v*_*t*_ versus *f*_*t*_

To examine if the long time between time points accounts for the perceived non-neutrality from our analysis of figure 2, we produced surrogate data by simulating neutral communities with similarly sparse time points. Parameter estimation of λ, *ρ* and WFP simulation is described in the supplement section S4. For one particular CVT, we simulated 4,000 independent trajectories to allow the superposition of the *v*_*t*_ vs *f*_*t*_ scatter plots from the male tongue data over the points from the surrogate data. Much of the strong mean reversion in the data can in fact be accounted for by the sparsity of time points (figure 4A), but the P-value distribution from constant-volatility tests of 16,000 randomly drawn surrogate CVT simulations is much more uniform than the same distribution from the male tongue, which has many small P-values indicative of consistent state-dependent volatility of *f*_*t*_ (figure 3b). Thus, the non-neutrality of the male tongue dataset is not due to long sampling intervals.

The male tongue microbiome deviates from neutrality by having a significantly more *β*_2_ > 0 than *β*_2_ < 0 for those auxiliary regressions yielding significant heteroskedasticity (figure 3C). *β*_2_ > 0 indicates that the volatility increases farther away from the mean and plotting the residuals, 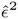 against the state variable, *f*_*t*_, reveals the form of heteroskedasticity (figure 3D). A similar significant hyper-abundance of *β*_2_ > 0 exists for all datasets considered here (figure 3E).

## Discussion

Evolution, driven by competition over rivalrous goods or limiting resources, is a phenomenon common to ecology, economics and sociology, and accurate statistical models of how competitive systems evolve can allow us to forecast, manage, and invest in them [2, 3, 8, 1, 31]. Neutral Theory is a null model of competition which assumes that all players are equal - that a canopy tree fills a gap in the canopy independent of its species’ identity, a dollar finds its way to another dollar independent of who owns the dollar, and a seat in congress is filled by someone independent of the racial, cultural or political traits of the successor or predecessor. It’s been hypothesized that neutrality could arise naturally as a result of competitively inferior species going extinct [32], and thus systems would tend towards neutrality over long periods of time, but the accuracy and generality of Neutral Theory as a dynamical model for a range of competitive systems was unclear.

We have provided a time-series test of neutral covariance structure that reveals a common feature of non-neutrality across a range of ecological and economic systems. Our test is based on the grouping invariance of neutral communities, and this grouping invariance is maintained by a particular co-variance structure of volatilities, namely where the volatility of a group, 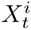, is 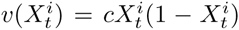 for some constant, *c*, which can only be invariant to grouping if the covariance between groups *i*, and *j* is 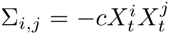. We test neutrality by randomly grouping species and testing if the volatility of those random groups is of the form of *v*(*x*). A deviation from *v* can indicate non-neutral covariance structure. If, for instance, instead of a quadratic curve of *v*, the volatility of random groups in the data follows a bell-shaped curve with positive curvature at the end-points, it could indicate that there is a positive covariance between rare species, possibly due to kill-the-winner effects or due to the relative abundances being driven by fluctuations in the largest populations. Conversely, excessive negative curvature could indicate strong negative covariances between rare species, possibly due to relatively constant populations of abundant species. Future work analyzing the volatility surfaces of random groups can improve our test and allow researchers to quickly isolate particular forms and fingerprints of common, non-neutral competitive asymmetries.

The results presented here are limited to the particular choice of species (the taxonomic scale), resource (trophic scale), and time scale. However, an analysis restricted to a particular scale means that our test can also allow researchers to probe multiple taxonomic, trophic and time-scales to see if there are patterns in which scales are most/least neutral in their dynamics. An alternative grouping of these original OTUs by genera may reveal different results by conditioning the groupings of species on a particular sub-set of possible groups, namely by grouping species with recent ancestors and shared traits together, and consequently this test could serve as a tool for evaluating competition at multiple taxonomic scales. The results are also limited to the choice of resource: companies within a sector governed by trust-busting policies which break the neutral symmetry in their market capitalization dynamics may still be neutral in their competition over the ethnic or cultural composition of their labor force. It’s possible that microbes in the gut are not neutral in their short-term fluctuations over the course of a year, but perhaps are neutral over longer time-scales that average out short-term fluctuations in diet and physiological state that are known to have predictable effects on microbial communities [33, 34].

Our test can be applied to any community of competing agents classified into discrete groups for which time-series of relative abundances (or market share, etc) of the groups are available. This test is most effective when the time-series is long and the spacing between samples is short relative to the turnover rate of the underlying resources (trees, dollars, congressional seats). There are many ways to build on our method. Explicit calculations of the drift and volatility over long time-intervals can improve our method for datasets with sparse time points. The dependence of the CVTs may be calculated as copulas allowing the implementation of a more exact goodness of fit test [35]. There may be more CVTs that solve the Hamilton-Jacobi equation, and different CVTs might be more specialized at detecting different asymmetries in competitive systems.

Neutrality tests of time-series data can help us understand the stochastic time-evolution of competitive systems and facilitate better prediction and management. For bacteria in the gut, for instance, understanding the important non-neutral forces governing the dynamics could allow progress towards the stochastic pharmacokinetics of probiotics [36]. Understanding the predictability of invasions at different taxonomic scales can tell us whether to evaluate metagenomic or ecological datasets based on the species, or whether other taxonomic levels will yield a more informative analysis - perhaps grouping tropical trees into the family Fabaceae, the family Melastomataceae, the genus *Cecropia*, and all other trees reveals trophic structure of tropical forests and competitive asymmetries that are drowned out by analyses at the species level. Demonstrating that the non-neutrality of portfolios is consistent with rare-species advantages in the Atlas model [8] would have major implications for portfolio design. In all cases, the first step of empirically demonstrating the existence of competitive asymmetries in time-series data can now be done with the test provided here.

## Materials and Methods

### Neutral Theory and the Wright-Fisher Process

Large neutral communities are well-approximated by a Wright-Fisher Process (WFP) [37, 38]. The convergence of discrete neutral communities to the continuous diffusion model of the WFP is covered in [39], and some numerical methods used for parameter estimation and simulation have been produced by [31]. The WFP is a continuous-state, continuous-time approximation of Kimura’s theory [14], it is an approximation of Hubbell’s neutral theory [15] when speciation rates are negligible over the timescale of interest, and it describes the dynamics of relative abundances of non-zero-sum volatility-stabilized market models [17, 27]. Using the WFP as a continuous approximation of large, finite communities simplifies the covariance in the jumps between species’ relative abundances, thereby permitting the analysis below.

The WFP models the stochastic time-evolution of relative abundances of *n* species. Let 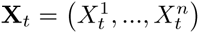 be the vector of relative abundances, the WFP is defined by the Ito SDE

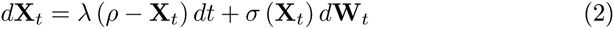

where λ > 0 and *ρ* > 0. The covariation between relative abundances of different species is given by the elements of Σ = σσ^*T*^/2, where

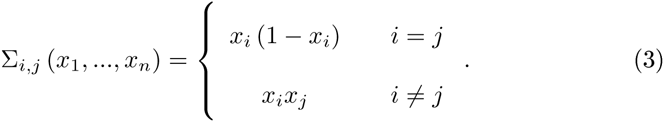

The deterministic motion, or drift, of the WFP - λ(*ρ* - X_*t*_) - yields exponential mean reversion like many dynamical systems reverting to a nearby stable equilibrium. The quadratic covariation of the WFP, Σ, captures the stochastic fingerprint of neutrality; it arises from randomly drawing a resource to be freed from its agent followed by randomly drawing one of the remaining agents to acquire that resource with a probability proportional to the agent’s current resource ownership. The family of Wright-Fisher Processes is closed to grouping, meaning that if a multi-species community’s dynamics are governed by a WFP, species can be grouped (e.g. collecting species into genera or higher taxonomic levels) and the dynamics of the resulting, re-grouped community will also be governed by a WFP.

### Testing Neutral Covariance Structure

We developed a test that is intentionally sensitive to the state-dependent noise of the WFP, allowing researches to test the underlying stochastic model of the random turnover of resources at the heart of neutral competition. Developing a strong test of the state-dependent covariance is not trivial, though, because direct measurement of the covariance of jumps conditioned on the state of the system, Cov [ΔX_*t*_|X_*t*_], would require many replicate time points each with the same initial state, X_*t*_, and, even with multiple time points at the same state, the sparsity of the high-dimensional data challenges the accurate estimation and significance testing of the covariance matrix. To circumvent these problems of replicate time points and high dimensionality and develop a strong test the state-dependent noise of the WFP, we find a variance-stabilizing transformation for the WFP that allows a regression-based heteroskedasticity test [40].

To be precise, we are looking for a real-valued function *f*(**X**_*t*_) such that for **X**_*t*_ obeying the WFP law in equation (1),

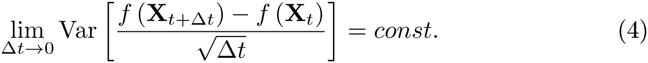

This approach is conceptually similar to the variance-stabilizing tools used for population fluctuation analyses [10, 25, 11] which stabilize the variance in jumps of a population size, except our function must stabilize the covariance of jumps between populations, not just the variance. In particular, to have a constant volatility, our function f must satisfy the Hamilton-Jacobi equation,

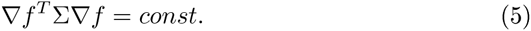

(see supplement part S2 for more details). The grouping invariance of the WFP can be used to intuit and show that there are at least 2^*n*^ different variance-stabilizing transformations of the WFP, parametrized by a vector a:

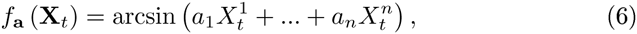

where *a*_*i*_ = ±1 for all *i*.

After transforming the data with *f*_*a*_, we need to perform a constant-volatility test. To test the constant volatility of *f*, we test the homoskedasticity of standardized jumps,

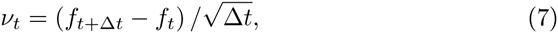

following regression on *f*_*t*_ to eliminate the state-dependent drift.

A homoskedasticity test of *v*_*t*_ for a single CVT is a test of neutrality for time-series data. However, with 2^*n*^ different CVTs, one can perform multiple hypothesis tests. For any multiple-hypothesis tests, if the null hypothesis is true, the distribution of P-values is uniform. In this paper, we test the uniformity of the distribution of resultant P-values from homoskedasticity tests of *v*_*t*_ for a number of randomly drawn CVTs. Figure 1 illustrates this test for 2,000 randomly drawn CVTs and shows the successful rejection of the WFP for the relative abundances of a mean-reverting geometric Brownian motion.

The P-values arising from homoskedasticity tests of different CVTs are not independent. Consequently, a Kolmogorov-Smirnov test of the P-value distribution against a uniform distribution would have a high false-positive rate. To reduce the false-positive rate, we perform a perturbation analysis to generate conservative estimates of cutoffs for the KS statistic at 0.05 and 0.005 significance levels. Details of the sensitivity analysis are provided in the supplement part S3.

## Acknowledgments

The authors would like to thank R. Chisholm, S. Levin, S. Pacala, J. O’dwyer, and J. B. Socolar for their discussions and feedback. In particular, A.D.W. would like to thank J. B. Socolar for numerous excellent recommendations that have greatly improved this manuscript, and both A. Gammie and D. Nemergut for their encouragement and support.

